# ΔFOSB in the nucleus accumbens core is required for increased anxiety, but not decreased social motivation, following estrogen withdrawal in female mice

**DOI:** 10.1101/2025.10.21.683778

**Authors:** Willow Clayton, May Y. Courtney, Lilah E. Craig, Helen E. Moniz, Alison B. Gibbons, Alissa A. Valentine, Achint K. Singh, Dylan R. Gearinger, Eric J. Nestler, Laura E. Been

## Abstract

During pregnancy, estrogen levels rise dramatically, but quickly drop to prepartum levels following birth, and remain suppressed until ovulation resumes. This “postpartum estrogen withdrawal” state has been linked to changes in the brain and behavior in humans and rodents. Previous research has demonstrated that following a hormone-simulated pseudopregnancy (HSP), an experimental model of postpartum estrogen withdrawal, female mice show increased anxiety-like behaviors and decreased social motivation. Further, these behavioral changes occur concurrently with an increase in ΔFOSB, a transcription factor associated with stable long-term plasticity, in the nucleus accumbens core. To test whether this increase in ΔFOSB is required for these behavioral changes, we used a viral-mediated gene transfer approach to prevent ΔFOSB-mediated transcription in the NAcC during HSP and found that it reduced the high-anxiety behavioral phenotype in estrogen-withdrawn females. However, preventing ΔFOSB-mediated transcription had little effect on social motivation. Together, these results suggest that postpartum estrogen withdrawal increases ΔFOSB in the NAc core to impact anxiety-like behaviors, but not social motivation, following estrogen withdrawal.

## Introduction

Circulating levels of estrogens increase significantly during pregnancy, creating a permissive environment for the growth and development of the fetus (Noyola-Martínez *et al*. 2019). After degeneration of the corpus luteum, estrogens are primarily secreted by the placenta (Albrecht & Pepe 2010), and increase nearly linearly across pregnancy, peaking at the end of the third trimester (Hendrick *et al*. 1998). Immediately following birth and the expulsion of the placenta, however, estrogen levels drop quickly and remain suppressed below pregravid levels until ovulation resumes (McNeilly 2001). The rapid transition from sustained high levels of estrogens during pregnancy to suppressed low levels estrogens during the early postpartum period has been hypothesized to create a “postpartum estrogen withdrawal” state, and has been linked to changes in mood and affect in humans (Sichel *et al*. 1995; Hendrick *et al*. 1998; Bloch 2000; Bloch *et al*. 2003; Douma *et al*. 2005).

A hormone simulated pseudopregnancy (HSP) has been used to model postpartum estrogen withdrawal in rodents. In this model, adult females are ovariectomized and then treated with daily hormone injections that model estrogen levels, and to a lesser extent progesterone levels, during pregnancy and the postpartum period. Initially developed in female Long-Evans rats (Galea *et al*. 2001), the model has been replicated (Green *et al*. 2009) and extended to Sprague-Daley rats (Stoffel & Craft 2004; Suda *et al*. 2008; Navarre *et al*. 2010; Schiller *et al*. 2013; Baka *et al*. 2017; Duan *et al*. 2025), hamsters (Hedges *et al*. 2021; Irvine *et al*. 2023), ICR mice (Zhang *et al*. 2016, 2017; Yang *et al*. 2017), and C57BL/6 mice (Buckhaults *et al*. 2023; Foster *et al*. 2023; Ren *et al*. 2024). Although the exact behavioral phenotypes vary by species and strain, withdrawal from estrogens after HSP is broadly associated with increased depressive-like and anxiety-like behaviors in rodents. In C57BL6 mice, postpartum estrogen withdrawal increases anxiety-like behavior in the elevated plus maze (Foster *et al*. 2023; Ren *et al*. 2024) and open field (Buckhaults *et al*. 2023; Ren *et al*. 2024), and decreases social motivation (Foster *et al*. 2023).

In addition to these changes in behavior, postpartum estrogen withdrawal following HSP has been associated with widespread neuroplastic changes in brain regions including the hippocampus (Suda *et al*. 2008; Green *et al*. 2009; Zhang *et al*. 2016, 2017; Baka *et al*. 2017; Ren *et al*. 2024), amygdala (Yang *et al*. 2017), hypothalamus (Hedges *et al*. 2021; Buckhaults *et al*. 2023; Irvine *et al*. 2023), bed nucleus of the stria terminalis (Buckhaults *et al*. 2023), dorsal raphe (Hedges *et al*. 2021), lateral habenula (Duan *et al*. 2025) and nucleus accumbens (NAc, Foster *et al*. 2023). Within the NAc, estrogen withdrawal following HSP results in an increase in ΔFOSB in both dopamine receptor 1 (D1) and dopamine receptor 2 (D2) neurons in the NAc Core (NAcC) of C57BL/6 mice (Foster *et al*. 2023). ΔFOSB is a transcription factor that regulates long-term neuroplastic changes following chronic or sustained stimulation (Nestler 2008). Interestingly, both ΔFOSB (Perrotti *et al*. 2004; Nestler 2015; Hamilton *et al*. 2018) and estrogen signaling (Lorsch *et al*. 2018; Georgiou *et al*. 2025) have been linked to transcriptional mechanisms underlying stress susceptibility. We therefore hypothesized that this increased ΔFOSB in the NAcC following estrogen withdrawal was causally related to behavioral changes in anxiety and motivation in the HSP model.

To test this hypothesis, we used a viral-mediated gene inhibition approach in combination with the HSP (Figure 1). Adult female mice were injected with an adeno-associated virus (AAV) vector expressing either ΔJUND or a control targeting the NAcC. ΔJUND is a dominant negative binding partner of ΔFOSB which, when virally expressed, competitively heterodimerizes and inhibits transcriptional effects of ΔFOSB (Berton *et al*. 2009; Been *et al*. 2013). We found that inhibiting ΔFOSB-mediated transcription reversed deficits in anxiety behaviors in estrogen-withdrawn animals in the elevated plus and open field but had minimal effect on social motivation. These data suggest that increased ΔFOSB in the NAcC following estrogen withdrawal is specifically and causally related to increased susceptibility to anxiety. These data may have important implications for the etiology and treatment of anxiety disorders during the postpartum period.

**Figure 1.**
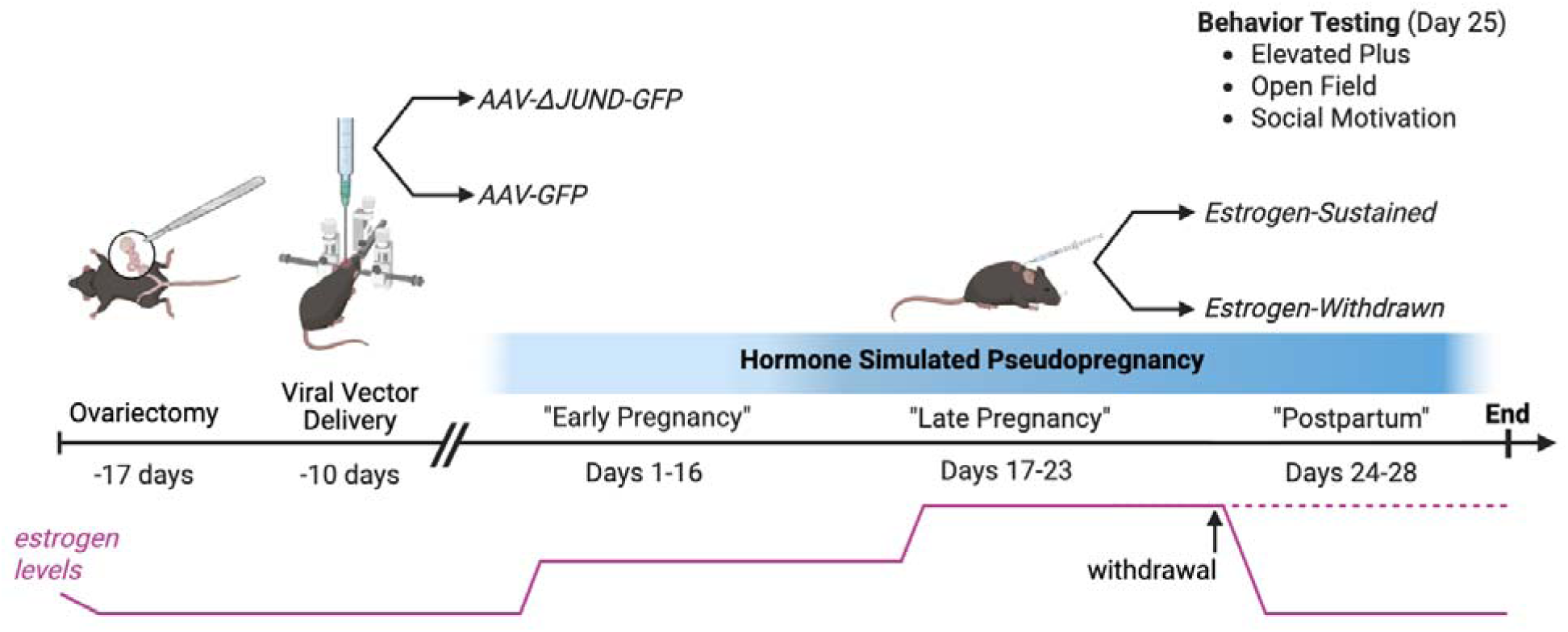
Experimental timeline. Adult, female C57BL/6 were ovariectomized and injected with an adeno-associated virus (AAV) vector containing either ΔJUND (AAV-ΔJUND-GFP) or a control vector (AAV-GFP) targeting the Nucleus Accumbens Core (NAcC). Subjects from both virus conditions began the hormone simulated pseudopregnancy regimen during which they received daily hormone injections approximating estrogen levels during pregnancy. After 23 days, half of the subjects were given vehicle injections for the duration of the “postpartum period,” simulating the rapid withdrawal of estrogen following birth. The other half of the subjects continued to receive high, late pregnancy-like levels of estrogen. All subjects underwent behavior testing on Day 25 and were euthanized on Day 28. Created in https://BioRender.com

## Material and Methods

### Subjects

Adult female C57BL/6 mice were ordered at 8-12 weeks of age from Charles Rivers Laboratories (Wilmington, MA, USA). Separate cohorts of animals were used for anxiety behavior testing (open field and elevated plus, n = 36) and social motivation testing (n = 28). Stimulus animals (n = 10) for social motivation tests were juvenile female C57BL/6 mice ordered at 3 weeks of age from Charles Rivers Laboratories. All mice were group-housed in plexiglass cages (28cm x 17cm x 12cm) with aspen bedding, cotton nesting material, and unlimited access to food and water. Animal holding rooms were temperature- and humidity-controlled and maintained on a 12h/12h reversed light-dark cycle (lights off at 8AM). All animal procedures were carried out in accordance with the National Institutes of Health (*Guide for the Care and Use of Laboratory Animals* 2011) and approved by the Haverford College Institutional Animal Care and Use Committee.

### Surgery

All surgery was conducted using aseptic surgical technique and under isoflurane anesthesia (2-3% vaporized in oxygen, Piramal, Bethlehem, PA, USA). EthiqaXR (1.3 mg/mL, Fidelis Animal Health, North Brunswick, NJ, USA) analgesic was administered subcutaneously immediately prior to the start of surgery.

#### Ovariectomy

Subjects’ bilateral flanks were shaved and cleaned with three alternating scrubs of 70% ethanol and Betadine before being transferred to a sterile surgical field. Isoflurane anesthesia was maintained via a nose cone. Subjects’ ovaries were extracted via bilateral incisions of the smooth muscle under the flanks and then removed via cauterization of the uterine horn and blood vessels. Polydioxanone absorbable suture (Ethicon, Sommerville, NJ, USA) and wound clips (Fine Science Tools, Foster City, CA, USA) were used to close the smooth muscle and skin incisions, respectively.

#### Viral Vector Injections

At least one week after ovariectomy, females received bilateral stereotaxic injections of AAV vectors targeting the NAcC. AAV is characterized by its ability to efficiently transfect neurons and maintain specific transgene expression for long periods of time (Chamberlin *et al*. 1998). This experiment used serotype 2 AAVs provided by the Nestler Laboratory with a titer of over 108/μl expressing constructs for either green fluorescent protein (GFP) and ΔJUND (*AAV-GFP-*Δ*JunD*), or a control AAV vector that contained only GFP (*AAV-GFP*). ΔJUND decreases ΔFOSB-mediated transcription by competitively heterodimerizing with ΔFOSB before binding the AP-1 region within gene promoters. These AAV vectors mediate transgene expression in mice that becomes maximal within 10 days of injection and then persists at this level for at least six months (Zachariou *et al*. 2006; Winstanley *et al*. 2007). Importantly, the vectors infect neurons only and produce no toxicity greater than vehicle infusions alone. Details for the production and use of these vectors are provided in earlier publications (Zachariou *et al*. 2006; Winstanley *et al*. 2007).

For *AAV-GFP-*Δ*JunD* or *AAV-GFP* vector delivery, subjects were anesthetized with isoflurane and secured in the stereotaxic apparatus such that their skull was level in the anterior-posterior (A-P) and medial-lateral (M-L) planes. Anesthesia was maintained via a nose cone. The dorsal scalp was shaved and cleaned with alternating scrubs of 70% ethanol and Betadine. Following a midline scalp incision, stereotaxic coordinates were measured relative to bregma (A-P: +1.4, M-L: ± 1.0). A hand-operated drill was used to expose dura. Stereotaxic injections were made by lowering a microinjection syringe (model number 7635-01, Hamilton, Reno, NV, USA) under stereotaxic control (Microinjection Unit, Model 5002, David Kopf Instruments, Tujunga, CA, USA) 4.5mm below dura and injecting 0.5 μl of AAV. To minimize the flow of AAV up the needle tract, the syringe was left in place for 5 min after each injection. The scalp incision was closed with wound clips. DietGel Recovery (Clear H_2_O, Portland, ME, USA) was provided in home cages for additional postoperative support.

### Hormone-Simulated Pseudopregnancy

Following recovery from stereotaxic surgery, subjects began the HSP injection regimen. On days 1-16, mice received daily subcutaneous injections containing a low dose (0.5 μg) of estradiol benzoate (Sigma) and a high dose (0.8 mg) of progesterone (Sigma) dissolved in 0.1mL cottonseed oil (Sigma). On days 17-23, mice received daily subcutaneous injections containing a high dose of estradiol benzoate (10 μg) in cottonseed oil. On days 24-28, mice were divided into two experimental conditions: estrogen-withdrawn (“withdrawn”) females received daily vehicle injections, modeling the dramatic drop in estrogen that occurs during the postpartum period, while females in the estrogen-sustained (“sustained”) group continued to receive daily injections of a high dose (10 μg) of estradiol. The doses and time course of the HSP hormone injections were initially chosen for their ability to elicit maternal responses in nulliparous, ovariectomized female rats (Siegel & Rosenblatt 1975; Rosenblatt 1988; Mayer *et al*. 1990), then modified for use in mice (Zhang *et al*. 2016). In rats, these doses produce serum concentrations that are supraphysiological for the high dose of estradiol, but are consistent with the level of progesterone reported during pregnancy (Bridges 1984; Garland *et al*. 1987; Galea *et al*. 2001).

### Behavior Testing

All behavior testing occurred during the dark phase. For cohort one, the order of testing (elevated plus vs. open field) was counterbalanced across virus and hormone conditions, with an inter-test interval of at least two hours.

#### Elevated Plus

On HSP day 25, subjects were tested for thigmotaxis-based anxiety in the elevated Plus. The apparatus consisted of a plus-shaped maze elevated 73 cm above the floor, with two open arms (51 x 11.5 cm) and two enclosed arms (51 x 11.5 x 39.5 cm). Animals were placed in the center of the maze facing a closed arm and allowed to freely explore the maze for five min under bright white light. Tests were recorded and the amount of time spent in each arm, as well as the total distance traveled and average velocity were quantified using Noldus Ethovision XT (version 14, Noldus Information Technology, Wageningen, The Netherlands).

#### Open Field

On HSP day 25, subjects were tested for thigmotaxis-based anxiety in the open field. The apparatus consisted of an open field (40.5 x 40.5 x 30cm) composed entirely of solid, light grey plastic (Maze Engineers, Glenview, IL, USA). Animals were placed in the center of the apparatus and allowed to freely explore for five min under bright white light. Tests were recorded and the amount of time spent in the center of the field, periphery of the field, as well as the total distance traveled and average velocity were quantified using Noldus Ethovision XT (version 14, Noldus Information Technology, Wageningen, The Netherlands).

#### Social Motivation Test

On HSP day 25, subjects were tested for social motivation using a two-choice preference test. The apparatus (Maze Engineers) consisted of a holding chamber with two removable slats that allowed for access to two larger chambers, one containing a social stimulus and one containing a nonsocial stimulus. The social stimulus chamber contained a juvenile female mouse under a mesh cup such that the experimental mouse could see, hear, and smell the stimulus mouse but could not contact it. The non-social stimulus chamber contained a black toy mouse of a similar shape, size, and color to the stimulus mouse (Penn-Plax play fur mouse toys, Hauppauge, NY, USA) under an identical mesh cup. The experimental mouse was acclimated to the closed holding chamber for 5 minutes before both slats were removed at which point the mouse could freely explore the entire testing apparatus. The social stimulus and non-social stimulus sides were alternated across subjects and the stimulus juvenile mouse used was changed after every trial. EthoVision XT software was used to track the movement of the mouse for ten minutes. The time spent on each side of the testing apparatus, in proximity to each stimulus, and actively investigating each stimulus was measured, along with average velocity and distance traveled.

### Histology

#### Immunohistochemistry

On day 26, subjects were anesthetized with isoflurane and rapidly decapitated. Brains were immediately removed and post-fixed in 4% paraformaldehyde overnight (4° C) and then cryoprotected for at least 48 hr in 30% sucrose in PBS. Coronal sections (35-μm) of brain tissue were sectioned on a cryostat (−20°C), collected in a 1:4 series, and stored at −20°C in cryoprotectant until immunohistochemical processing.

Sections containing the NAc were stained for GFP to confirm injection placement and expression. Tissue was washed 5 x 5 minutes in 25 mM PBS, then incubated in a primary antibody against GFP (Goat anti-GFP, 1:500, Abcam, Cambridge, UK) in PBS with 0.1% Triton-X (PBTX) for 48 hours. Following primary antibody incubation, the tissue again underwent 5 x 5 minute PBS washes before being incubated in fluorescent secondary antibody (anti-goat Alexa488, 1:500, Jackson Immunoresearch Lab, WestGrove, PA, USA) in 0.1% PBTx for one hour. Finally, tissue was washed 5 x 5 minutes in PBS, mounted onto slides, and coverslipped with Prolong Gold mounting media (product info) for microscopy.

#### Microscopy

An epifluorescent microscope (Eclipse 400, Nikon, Melville, NY, USA) was used to visualize the viral injections tissue and acquire photomicrograph images. Images were collected at a 10x magnification of the NAc via NIS Elements software (Nikon). Animals with incomplete viral transfection or injection placement outside of the NAcC were excluded from analysis.

### Statistical Analysis

For the elevated plus, thigmotaxis difference scores were calculated to determine the amount of time spent in the open vs. closed arms of the apparatus (time spent in the open arm - time spent in closed arm). For the open field, thigmotaxis difference scores were calculated to determine the amount of time spent in the periphery vs. the center of the apparatus (time spent in periphery - time spent in center). Difference scores were analyzed using 2 x 2 factorial ANOVAs to probe for significant interactions and/or main effects of virus condition (JUND vs. GFP) and hormone condition (sustained vs. withdrawn) on behavior. For both behavior tests, distance traveled and average velocity were also compared across groups using 2 x 2 factorial ANOVAs. Significant effects were explored with post hoc tests using Bonferroni corrections for multiple comparisons.

For the social motivation test, 2 x 2 factorial ANOVAs were used to probe for significant interactions and/or main effects of virus condition (JUND vs. GFP) and hormone condition (sustained vs. withdrawn) on social investigation preference (social stimulus investigation duration - non-social stimulus investigation duration). In addition, 2 x 2 factorial ANOVAS were used to probe for significant interactions and/or main effects of virus condition (JUND vs. GFP) and hormone condition (sustained vs. withdrawn) on time spent on the side of the chamber containing the social stimulus, time spent in the side of the chamber containing the non-social stimulus, time spent in proximity to the social stimulus, and time spent in proximity to the non-social stimulus. Significant effects were explored with post hoc tests using Bonferroni corrections for multiple comparisons.

## Results

### Injection Placement

In the anxiety behavior cohort, 2 mice did not recover from surgery and one mouse died during the HSP protocol. Of the remaining 33 mice, 4 mice were removed from the study due to incomplete or misplaced injections (JUND-withdrawn n = 1, GFP-withdrawn n = 2, JUND-sustained n = 1), bringing the total number of mice included in the analyses to n = 29. In the social motivation cohort, two mice did not recover from surgery and 2 animals were removed from the study due to incomplete or misplaced injections (JUND-withdrawn n = 1, JUND-sustained n = 1), bringing the total number of mice included in the analyses to n = 24. All remaining subjects had significant viral expression in the NAcC, and minimal spread to adjacent brain regions (see Figure 2).

**Fig 2:**
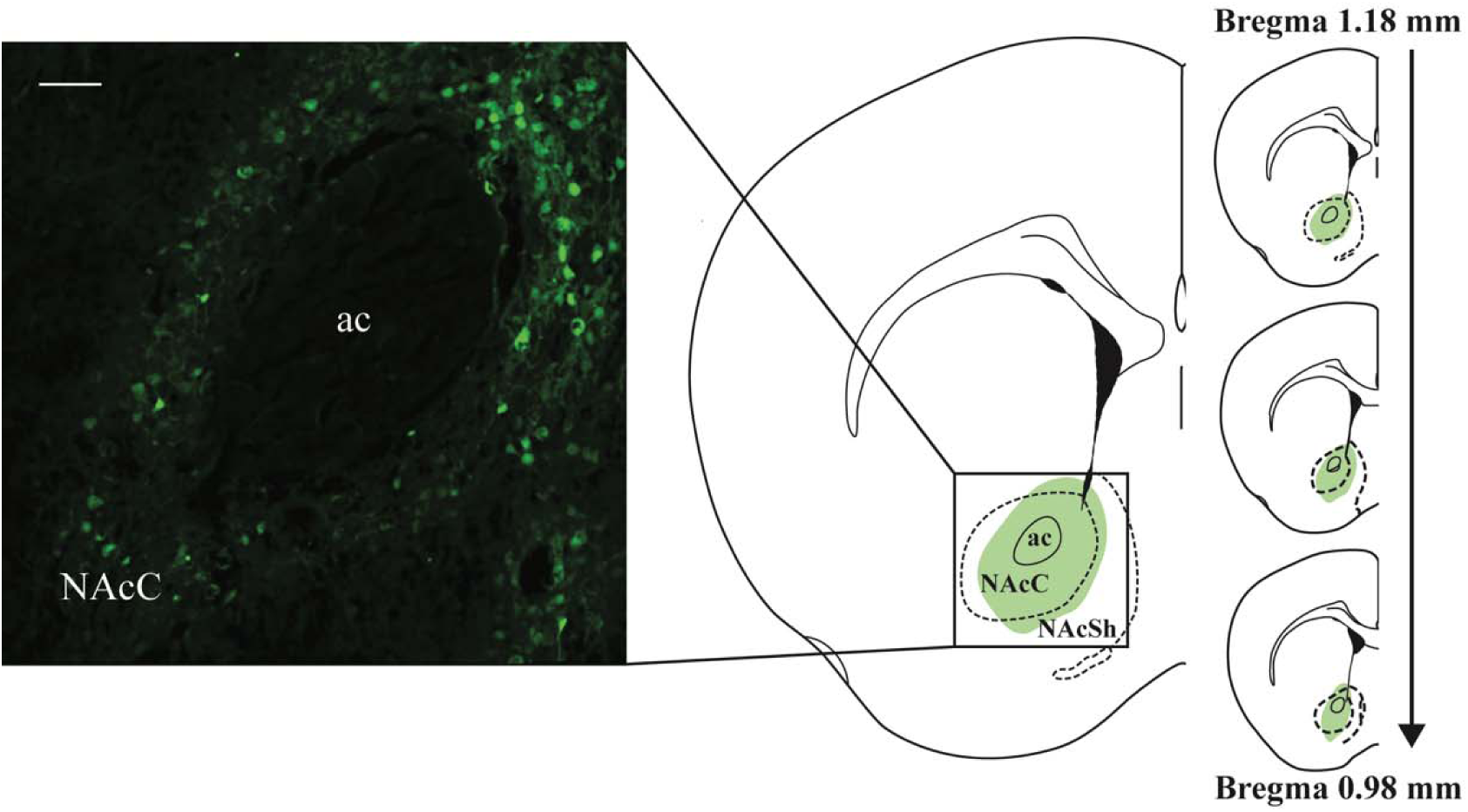
Validation of viral Injections. Sections containing the NAc were stained for GFP to confirm injection placement and expression. Immunostaining revealed robust transfection including cell bodies and processes (see representative photomicrograph, left; scale bar = 100 μm). Valid injection placements surrounded the anterior commissure (ac) and transfected the majority the NAcC. Some larger injections partially transfected the NAcSh. Injection sites typically spanned 0.2 mm in the mid-caudal NAc. Animals with incomplete viral transfection of the NAcC or injection placement significantly outside of the NAcC were excluded from analysis. Atlas plates modified from The Mouse Brain in Stereotaxic Coordinates (Paxinos & Franklin 2019).

**Fig 3:**
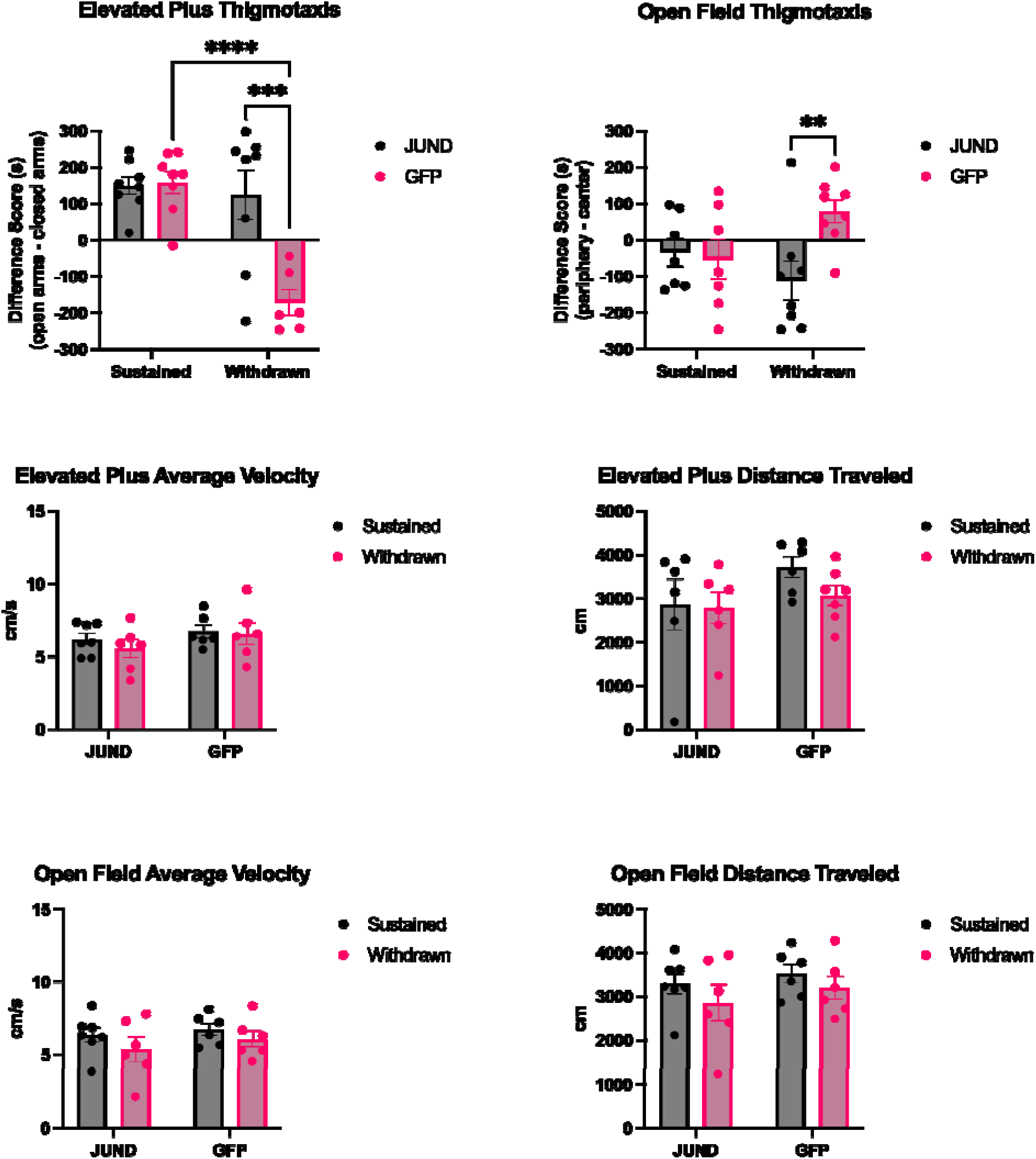
Preventing ΔFOSB-mediated transcription in the NAcC reverses the anxiogenic effect of estrogen-withdrawal following HSP. In the elevated plus, there was a significant interaction between virus condition and hormone condition on the thigmotaxis difference score (time spent in the open arm - time spent in closed arm). Estrogen-withdrawn mice who received the control *AAV-GFP vector* (GFP-withdrawn) spent more time in closed arm, indicating higher anxiety. However, anxiety associated with estrogen-withdrawal was prevented in subjects that received the ΔJun-AAV-GFP vector (JUND-withdrawn), which prevents ΔFOSB-mediated transcription. Likewise in the open field, there was a significant interaction between virus condition and hormone condition on the thigmotaxis difference score (time spent in periphery - time spent in center). GFP-withdrawn mice spent more time in the periphery, indicating higher anxiety. However, anxiety associated with estrogen-withdrawal was prevented in JUND-withdrawn mice who spent significantly more time in the center of the open field than GFP-withdrawn mice. General locomotor behaviors (average velocity or distance traveled) did not differ across hormone or virus conditions. **p < 0.01, *** p < 0.001, ****p < 0.0001.

**Fig 4:**
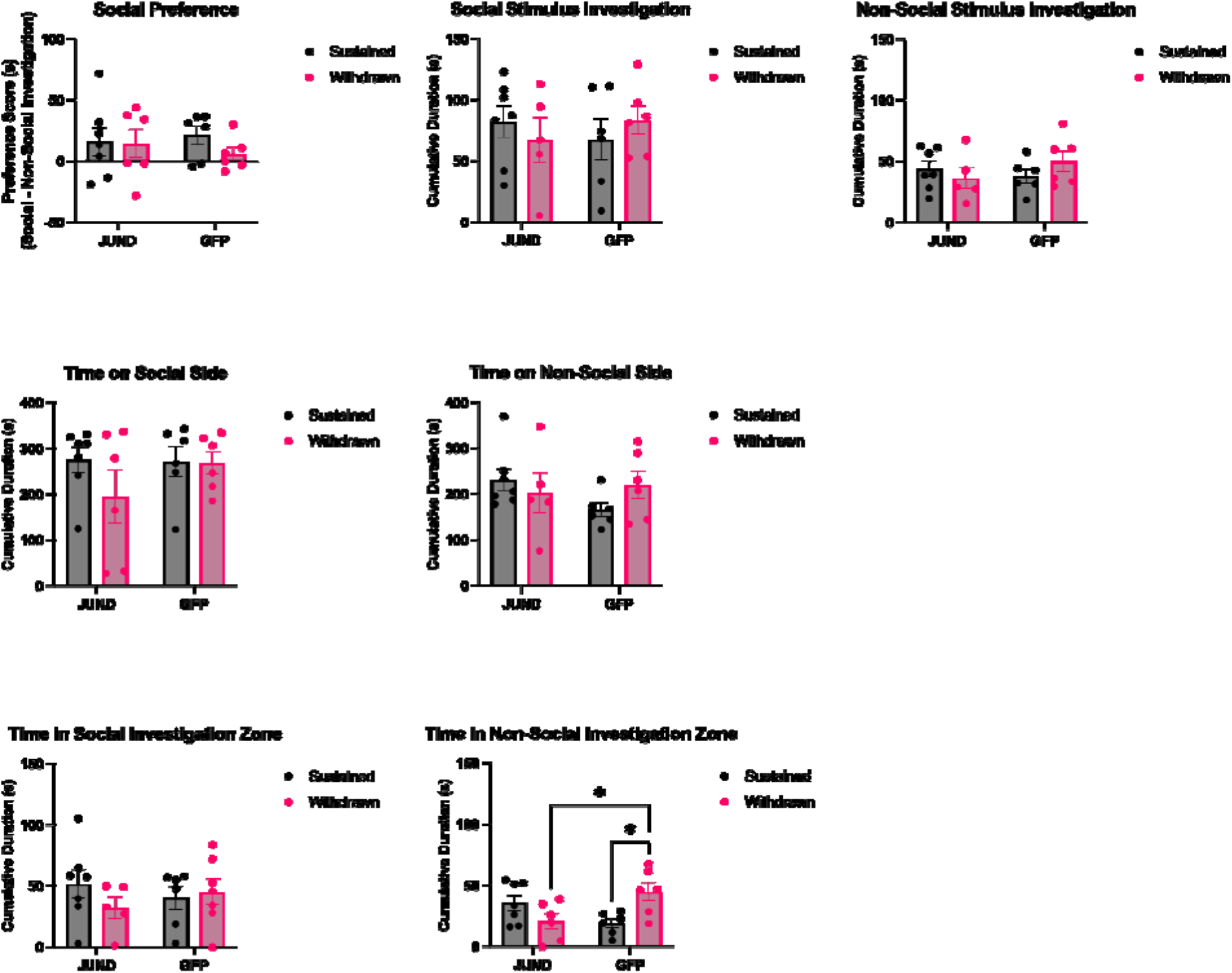
Preventing ΔFOSB-mediated transcription in the NAcC does not impact social motivation following HSP. In the social motivation assay, most animals preferred to investigate the social stimulus over the non-social stimulus regardless of virus or hormone condition. There was no effect of either hormone condition or virus condition on social preference, social stimulus investigation, non-social stimulus investigation, the time spent on the social side of the apparatus, the time spend on the non-social side of the apparatus, or the time spent in the social investigation zone. However, there was an interaction between virus condition and hormone condition on the amount of time spent in proximity to the non-social stimulus. GFP-withdrawn mice spent significantly more time in proximity to the non-social stimulus than GFP-sustained mice, whereas JUND-withdrawn animals spent significantly less time in proximity to the non-social stimulus than GFP-withdrawn animals. *p < 0.05

### Elevated Plus

In the elevated plus maze, thigmotaxis-based anxiety measures varied with both virus condition and hormone condition. Specifically, there was a significant interaction between virus condition and hormone condition on the thigmotaxis difference score (F(1, 26) = 11.75, p = 0.0020). Post-hoc tests revealed that JUND-withdrawn mice spent significantly more time in the open arm than GFP-withdrawn mice (p < 0.0001). Further, GFP-sustained mice spent significantly more time in the open arm than GFP-withdrawn mice (p < 0.0001). There was no difference in difference scores between JUND-sustained and GFP-sustained mice, or between JUND-sustained and JUND-withdrawn mice (both p < 0.05).

General locomotor behaviors did not differ across hormone or virus conditions. Specifically, there was no significant interaction between virus condition and hormone condition on average velocity in the elevated plus (F(1, 21) = 0.2073, p = 0.6536), nor was there a main effect of virus condition (F(1, 21) = 1.831, p = 0.1904) or hormone condition (F(1, 21) = 0.5003, p = 0.4872). There was also no significant interaction between virus condition and hormone condition on distance traveled in the elevated plus (F(1, 21) = 0.5625, p = 0.4616), nor was there a main effect of virus condition (F(1, 21) = 2.311, p = 0.1417) or hormone condition (F(1, 21) = 0.9390, p = 0.3436).

### Open Field

In the open field, thigmotaxis-based anxiety measures varied with both virus condition and hormone condition. Specifically, there was a significant interaction between virus condition and hormone condition on the thigmotaxis difference score (F(1, 26) = 5.318, p = 0.0293). Post-hoc tests revealed JUND-withdrawn mice spent significantly more time in the center of the open field than GFP-withdrawn mice (p = 0.0050). Although there was a trend for GFP-withdrawn mice to spend more time in the center of the open field than GFP-sustained mice, this difference did not reach significance (p = 0.0503). Further, there was no significant difference in thigmotaxis difference score between JUND-sustained vs. JUND-withdrawn mice, or between JUND-sustained vs. GFP-sustained mice (all p < 0.05).

General locomotor behaviors did not differ across hormone or virus conditions. Specifically, there was no significant interaction between virus condition and hormone condition on average velocity in the open field (F(1, 21) = 0.07889, p = 0.7816), nor was there a main effect of virus condition (F(1, 21) = 0.7981, p = 0.3818) or hormone condition (F(1, 21) = 1.1775, p = 0.1970). There was also no significant interaction between virus condition and hormone condition on distance traveled in the open Field F(1, 21) = 0.03123, p = 0.8614), nor was there a main effect of virus condition (F(1, 21) = 0.9673, p = 0.3365) or hormone condition (F(1, 21) = 1.675, p = 0.2096).

### Social Motivation

In the social motivation assay, most animals preferred to investigate the social stimulus over the non-social stimulus regardless of virus or hormone condition. Although withdrawn animals spent less time on average investigating the social stimulus, there was no interaction between virus condition and hormone condition on social investigation preference (F(1, 21) = 0.4586, p = 0.5057), nor was there a main effect of virus condition (F(1, 21) = 0.0219, p = 0.8837) or hormone condition (F(1, 21) = 0.7216, p = 0.4052). In addition, there was no significant interaction between virus condition and hormone condition on the time spent on the side of the chamber that contained the social stimulus (F(1, 21) = 1.032, p = 0.3212), nor was there a main effect of virus condition (F(1, 21) = 0.8673, p = 0.3623) or hormone condition (F (1, 21) = 1.200, p = 0.2858). Likewise, there was no significant interaction between virus condition and hormone condition on the time spent on the side of the chamber that contained the non-social stimulus (F(1, 21) = 2.051, p = 0.1676), nor was there a main effect of virus condition (F(1, 21) = 0.6998, p = 0.4127) or hormone condition (F(1,21) = 0.2136, p = 0.6489). Finally, there was no significant interaction between virus condition and hormone condition on the amount of time spent in proximity to the social stimulus (F(1, 21) = 1.289, p = 0.2691, nor was there a main effect of virus condition (F(1,21) = 0.004461, p = 0.9474) or hormone condition (F(1, 21) = 0.4460, p = 0.5115). However, there was an interaction between virus condition and hormone condition on the amount of time spent in proximity to the non-social stimulus (F(1, 21) = 10.51, p = 0.0039). Post-hoc tests revealed that GFP-withdrawn mice spent significantly more time in proximity to the non-social stimulus (p = 0.0096) than GFP-sustained mice, whereas JUND-withdrawn animals spent significantly less time in proximity to the non-social stimulus than GFP-withdrawn animals.

## Discussion

Here we demonstrate for the first time that increased ΔFOSB in the NAcC is causally linked to increased anxiety-like behavior in a mouse model of postpartum estrogen withdrawal. Specifically, preventing ΔFOSB-mediated transcription in the NAcC during HSP through viral expression of its dominant negative antagonist, ΔJUND, prevents the increase in anxiety-like behavior typically seen in estrogen-withdrawn females. Interestingly, this change appears to be specific to anxiety-like behaviors as manipulating ΔFOSB in the NAcC generally did not impact social motivation. This suggests that ΔFOSB-mediated transcription in the NAcC is necessary to confer long-term plasticity required for anxiety-like behavioral changes following postpartum estrogen withdrawal. In the current study, expression of ΔJUND was targeted to the NAcC with minimal spread to the NAc shell or other brain areas outside of the NAc. The ability of core-centered injections of ΔJUND to alter anxiety-like behaviors following HSP is in agreement with previous research from our laboratory demonstrating that estrogen withdrawal following HSP is associated with increased numbers of FOSB-immunoreactive cells in the NAcC, but not the NAc shell, as well as increased ΔFOSB protein in NAcC-centered tissue punches (Foster *et al*. 2023).

Estrogen withdrawal following HSP has been shown to increase anxiety-like behaviors in a range of rodent species, although there is variability across different species, strains, and behavioral tests. The majority of studies in mice, including ICR mice (Zhang *et al*. 2016; Yang *et al*. 2017) and C57BL/6 mice (Foster *et al*. 2023; Ren *et al*. 2024), find increases in thigmotaxis-based anxiety measures in the elevated plus, although a recent study in C57BL6 mice did not replicate this finding (Buckhaults *et al*. 2023). Likewise, the majority of studies in mice also find increases in thigmotaxis-based anxiety measures in the open field (Zhang *et al*. 2016, 2017; Yang *et al*. 2017; Buckhaults *et al*. 2023), although this difference did not reach significance in a recent study in C57BL6 mice (Foster *et al*. 2023). In the current study, estrogen withdrawal following HSP resulted in increased anxiety-like behavior in both the elevated plus and open field, and these behavioral changes were prevented by blocking ΔFOSB-mediated transcription in NAcC.

Increased accumulation of ΔFOSB in the NAc is typically correlated with chronic stimulation of the mesolimbic pathway, which is hypothesized to increase the sensitivity of the NAc to future rewarding stimuli. This has been best studied following chronic exposure to drugs of abuse, which induce high levels of ΔFOSB in the NAc and lead to long-lasting behavioral responses that are indicative of an addiction phenotype. However, there are moderate levels of ΔFOSB in the NAc under baseline conditions, and these levels are modified by exposure to both rewarding and aversive stimuli over long periods of time (Nestler *et al*. 2001; Nestler 2008). The induction of ΔFOSB by endogenous stimuli may therefore facilitate adaptive responding to environmental challenges. Chronic high levels of estrogens during pregnancy may act on the estrogen-sensitive mesolimbic pathway, altering DA dynamics and increasing ΔFOSB levels in the NAc. In the HSP model, these increased levels of ΔFOSB not only persist, but are enhanced, following the withdrawal of estrogen. This is consistent with the protein stability of ΔFOSB even in the absence of further stimulation. The concurrent increase in anxiety-like behaviors following estrogen withdrawal may therefore reflect an adaptive response to the new demands of parenthood and may be more accurately framed as an increase in vigilance.

Given the known relationship between ΔFOSB and motivation, we hypothesized that blocking ΔFOSB-mediated transcription in NAcC would prevent the deficits in social motivation observed in estrogen-withdrawn mice. However, in the current study, expression of ΔJUND in the NAcC had minimal effect on social motivation. The most straightforward interpretation of these findings is that ΔFOSB accumulation in the NAcC does not underlie deficits in social motivation following estrogen withdrawal. However, it is worth considering how the parameters of the social motivation assay may influence this interpretation. Specifically, the social stimulus used in the current assay was a novel juvenile female mouse. This stimulus was chosen because it was unlikely to elicit an aggressive or sexual response, and approach behavior would therefore reflect unconditioned social motivation. Indeed, previous studies in mice have shown that stimulation of VTA DA neurons increases social approach (Gunaydin *et al*. 2014), whereas inhibition of VTA DA neurons attenuates social exploration of a nonfamiliar conspecific (Bariselli *et al*. 2018). However, recent work suggests that DA release during social investigation may primarily signal the novelty of the stimulus, as a novel mouse elicits higher DA responses than a familiar mouse during a simultaneous presentation preference test (Dai *et al*. 2022). Given that both the social and non-social stimuli in the current task were novel stimuli, it is possible that both stimuli activated enhanced mesolimbic signaling that result from chronic stimulation during the HSP. Future experiments may wish to use a familiar social stimulus (e.g., a cage mate) to overcome this limitation.

Previous studies using the HSP model have used sucrose preference as an additional readout of motivated behavior. Although the first studies in rats found a reduction in sucrose preference (Green *et al*. 2009), more recent studies in hamsters found no effect of HSP on sucrose preference (Hedges *et al*. 2021), and studies in C57BL6 mice found that estrogen withdrawal enhances sucrose preference (Buckhaults *et al*. 2023; Foster *et al*. 2023). In particular, the findings in mice are consistent with the interpretation that estrogen withdrawal following HSP alters DA dynamics in the NAcC. In fact, the increase in ΔFOSB in the NAcC in estrogen withdrawn animals may result from an increase in phasic DA release into the NAc. This interpretation is supported by the finding that hormone treatment during HSP increases not just sucrose preference, but also sucrose consumption, in C57BL6 mice (Foster *et al*. 2023). Further, phasic-like dopamine release is increased in naturally parturient female rats as measured by in vivo fast scan cyclic voltammetry (Shnitko *et al*. 2017). Ongoing experiments in our laboratory are testing the impact of estrogen withdrawal following HSP on tonic and phasic dopamine release in the NAcC of C57BL6 mice. Future experiments should also test the requirement of ΔFOSB for enhanced sucrose preference following HSP.

A remaining question is whether the effects of preventing ΔFOSB-mediated transcription in NAcC on anxiety-like behavior is mediated specifically by D1 or D2 neurons. The expression of ΔJUND in the current study does not selectively impact one neuronal subtype, but rather inhibits ΔFOSB-mediated transcription in all transfected neurons. Previous work in our lab has shown that estrogen withdrawal following HSP increases ΔFOSB in both D1 and D2 neurons. However, it remains possible that one neuronal subtype is more critical for the relationship between estrogen withdrawal, ΔFOSB, and anxiety-like behaviors. Indeed, susceptibility to social defeat stress is differentially regulated by FOSB gene products (presumably ΔFOSB) in D1 vs. D2 neurons (Hamilton *et al*. 2018). Given that ΔFOSB has been shown to increase not only during rewarding experiences, but also during chronic stress (Perrotti *et al*. 2004), future experiments could investigate the possibility of cell-type-specificity by injection Cre-dependent viral vectors in the NAc of transgenic mice that selectively express Cre recombinase in either D1 or D2 cell types following HSP. Nonetheless, the data presented here are the first to link ΔFOSB to anxiety-like behavior following estrogen withdrawal. This has important implications for understanding mechanisms of neuroplasticity that may lead to both adaptive and maladaptive behavioral responses during the early postpartum period.

## Acknowledgements

This work was funded by NIMH Grants R15MH125282 to LEB and R01MH129306 to EJN. The authors would like to thank Dr. Peter Hamilton (Virginia Commonwealth University) for coordinating the shipping of the viral vectors from Icahn School of Medicine to Haverford College.

## Notes

### Competing Interest Statement

The authors have declared no competing interest.

## References

Albrecht, E.D. & Pepe, G.J. (2010) Estrogen regulation of placental angiogenesis and fetal ovarian development during primate pregnancy. Int J Dev Biol 54, 397–408.

Baka, J., Csakvari, E., Huzian, O., Dobos, N., Siklos, L., Leranth, C., MacLusky, N.J., Duman, R.S. & Hajszan, T. (2017) Stress induces equivalent remodeling of hippocampal spine synapses in a simulated postpartum environment and in a female rat model of major depression. Neuroscience 343, 384–397.

Bariselli, S., Hörnberg, H., Prévost-Solié, C., Musardo, S., Hatstatt-Burklé, L., Scheiffele, P. & Bellone, C. (2018) Role of VTA dopamine neurons and neuroligin 3 in sociability traits related to nonfamiliar conspecific interaction. Nat Commun 9, 3173.

Been, L.E., Hedges, V.L., Vialou, V., Nestler, E.J. & Meisel, R.L. (2013) ΔJunD overexpression in the nucleus accumbens prevents sexual reward in female Syrian hamsters: ΔJunD prevents sexual reward. *Genes*, Brain and Behavior 12, 666–672.

Berton, O., Guigoni, C., Li, Q., Bioulac, B.H., Aubert, I., Gross, C.E., DiLeone, R.J., Nestler, E.J. & Bezard, E. (2009) Striatal Overexpression of ΔJunD Resets L-DOPA-Induced Dyskinesia in a Primate Model of Parkinson Disease. Biological Psychiatry 66, 554–561.

Bloch, M. (2000) Effects of Gonadal Steroids in Women With a History of Postpartum Depression. American Journal of Psychiatry 157, 924–930.

Bloch, M., Daly, R.C. & Rubinow, D.R. (2003) Endocrine factors in the etiology of postpartum depression. Comprehensive Psychiatry 44, 234–246.

Bridges, R.S. (1984) A Quantitative Analysis of the Roles of Dosage, Sequence, and Duration of Estradiol and Progesterone Exposure in the Regulation of Maternal Behavior in the Rat *. Endocrinology 114, 930–940.

Buckhaults, K., Swack, B.D. & Sachs, B.D. (2023) Estrogen administration and withdrawal in a model of hormone-simulated pregnancy lead to alterations in behavior and gene expression but do not induce depression-like phenotypes in mice. Physiology & Behavior 269, 114288.

Chamberlin, N.L., Du, B., De Lacalle, S. & Saper, C.B. (1998) Recombinant adeno-associated virus vector: use for transgene expression and anterograde tract tracing in the CNS. Brain Research 793, 169–175.

Dai, B., Sun, F., Tong, X., Ding, Y., Kuang, A., Osakada, T., Li, Y. & Lin, D. (2022) Responses and functions of dopamine in nucleus accumbens core during social behaviors. Cell Reports 40, 111246.

Douma, S.L., Husband, C., O Donnell, M.E., Barwin, B.N. & Woodend, A.K. (2005) Estrogen-related Mood Disorders: Reproductive Life Cycle Factors. Advances in Nursing Science 28, 364–375.

Duan, C., Ma, S., Chen, M., Wang, J., Jiang, Y., Ye, M., Tan, Y., Cheng, S., Yang, X., Hu, H., Yang, Y. & Huang, H.-F. (2025) Estrogen receptor beta in lateral habenula mediates antidepressant effects of estrogen in postpartum-hormone-withdrawal-induced depression. Mol Psychiatry.

Foster, W.B., Beach, K.F., Carson, P.F., Harris, K.C., Alonso, B.L., Costa, L.T., Simamora, R.C., Corbin, J.E., Hoag, K.F., Mercado, S.I., Bernhard, A.G., Leung, C.H., Nestler, E.J. & Been, L.E. (2023) Estradiol withdrawal following a hormone simulated pregnancy induces deficits in affective behaviors and increases ΔFosB in D1 and D2 neurons in the nucleus accumbens core in mice. Hormones and Behavior 149, 105312.

Galea, L.A.M., Wide, J.K. & Barr, A.M. (2001) Estradiol alleviates depressive-like symptoms in a novel animal model of post-partum depression. Behavioural Brain Research 122, 1–9.

Garland, H.O., Atherton, J.C., Baylis, C., Morgan, M.R.A. & Milne, C.M. (1987) Hormone profiles for progesterone, oestradiol, prolactin, plasma renin activity, aldosterone and corticosterone during pregnancy and pseudopregnancy in two strains of rat: correlation with renal studies. Journal of Endocrinology 113, 435– 444.

Georgiou, P., Postle, A.F., Mou, T.-C.M., Potter, L.E., An, X., Zanos, P., Patton, M.S., Pultorak, K.J., Clark, S.M., Ngyuyen, V., Powels, C.F., Prokai-Tatrai, K., Kirmizis, A., Merchenthaler, I., Prokai, L., McCarthy, M.M., Mathur, B.N. & Gould, T.D. (2025) Estradiol, via estrogen receptor β signaling, mediates stress-susceptibility in the male brain. Mol Psychiatry 30, 4445–4459.

Green, A.D., Barr, A.M. & Galea, L.A.M. (2009) Role of estradiol withdrawal in “anhedonic” sucrose consumption: a model of postpartum depression. Physiol Behav 97, 259–265.

Guide for the Care and Use of Laboratory Animals: Eighth Edition. (2011). National Academies Press, Washington, D.C.

Gunaydin, L.A., Grosenick, L., Finkelstein, J.C., Kauvar, I.V., Fenno, L.E., Adhikari, A., Lammel, S., Mirzabekov, J.J., Airan, R.D., Zalocusky, K.A., Tye, K.M., Anikeeva, P., Malenka, R.C. & Deisseroth, K. (2014) Natural Neural Projection Dynamics Underlying Social Behavior. Cell 157, 1535–1551.

Hamilton, P.J., Burek, D.J., Lombroso, S.I., Neve, R.L., Robison, A.J., Nestler, E.J. & Heller, E.A. (2018) Cell-Type-Specific Epigenetic Editing at the Fosb Gene Controls Susceptibility to Social Defeat Stress. Neuropsychopharmacology: Official Publication of the American College of Neuropsychopharmacology 43, 272–284.

Hedges, V.L., Heaton, E.C., Amaral, C., Benedetto, L.E., Bodie, C.L., D’Antonio, B.I., Davila Portillo, D.R., Lee, R.H., Levine, M.T., O’Sullivan, E.C., Pisch, N.P., Taveras, S., Wild, H.R., Grieb, Z.A., Ross, A.P., Albers, H.E. & Been, L.E. (2021) Estrogen Withdrawal Increases Postpartum Anxiety via Oxytocin Plasticity in the Paraventricular Hypothalamus and Dorsal Raphe Nucleus. Biol Psychiatry 89, 929–938.

Hendrick, V., Altshuler, L.L. & Suri, R. (1998) Hormonal changes in the postpartum and implications for postpartum depression. Psychosomatics 39, 93–101.

Irvine, A., Gaffney, M.I., Haughee, E.K., Horton, M.A., Morris, H.C., Harris, K.C., Corbin, J.E., Merrill, C., Perlis, M.L. & Been, L.E. (2023) Elevated estradiol during a hormone simulated pseudopregnancy decreases sleep and increases hypothalamic activation in female Syrian hamsters. J Neuroendocrinology 35, e13278.

Lorsch, Z.S., Loh, Y.-H.E., Purushothaman, I., Walker, D.M., Parise, E.M., Salery, M., Cahill, M.E., Hodes, G.E., Pfau, M.L., Kronman, H., Hamilton, P.J., Issler, O., Labonté, B., Symonds, A.E., Zucker, M., Zhang, T.Y., Meaney, M.J., Russo, S.J., Shen, L., Bagot, R.C. & Nestler, E.J. (2018) Estrogen receptor α drives pro-resilient transcription in mouse models of depression. Nat Commun 9, 1116.

Mayer, A.D., Ahdieh, H.B. & Rosenblatt, J.S. (1990) Effects of prolonged estrogen-progesterone treatment and hypophysectomy on the stimulation of short-latency maternal behavior and aggression in female rats. Horm Behav 24, 152–173.

McNeilly, A.S. (2001) Lactational control of reproduction. Reprod Fertil Dev 13, 583– 590.

Navarre, B.M., Laggart, J.D. & Craft, R.M. (2010) Anhedonia in postpartum rats. Physiol Behav 99, 59–66.

Nestler, E.J. (2008) Transcriptional mechanisms of addiction: role of ΔFosB. Philos Trans R Soc Lond B Biol Sci 363, 3245–3255.

Nestler, E.J. (2015) ΔFosB: A transcriptional regulator of stress and antidepressant responses. European Journal of Pharmacology 753, 66–72.

Nestler, E.J., Barrot, M. & Self, D.W. (2001) DeltaFosB: a sustained molecular switch for addiction. Proc Natl Acad Sci U S A 98, 11042–11046.

Noyola-Martínez, N., Halhali, A. & Barrera, D. (2019) Steroid hormones and pregnancy. Gynecological Endocrinology 35, 376–384.

Paxinos, G. & Franklin, K.B.J. (2019) Paxinos and Franklin’s The mouse brain in stereotaxic coordinates. Fifth edition. Elsevier, Academic Press, London San Diego Cambridge; MA Kidlington, Oxford.

Perrotti, L.I., Hadeishi, Y., Ulery, P.G., Barrot, M., Monteggia, L., Duman, R.S. & Nestler, E.J. (2004) Induction of ΔFosB in Reward-Related Brain Structures after Chronic Stress. J Neurosci 24, 10594–10602.

Ren, P., Wang, J.-Y., Chen, H.-L., Wang, Y., Cui, L.-Y., Duan, J.-Y., Guo, W.-Z., Zhao, Y.-Q. & Li, Y.-F. (2024) Activation of σ-1 receptor mitigates estrogen withdrawal-induced anxiety/depressive-like behavior in mice via restoration of GABA/glutamate signaling and neuroplasticity in the hippocampus. Journal of Pharmacological Sciences 154, 236–245.

Rosenblatt, J. (1988) Hormonal basis during pregnancy for the onset of maternal behavior in the rat. Psychoneuroendocrinology 13, 29–46.

Schiller, C.E., O’Hara, M.W., Rubinow, D.R. & Johnson, A.K. (2013) Estradiol modulates anhedonia and behavioral despair in rats and negative affect in a subgroup of women at high risk for postpartum depression. Physiol Behav 119, 137–144.

Shnitko, T.A., Mace, K.D., Sullivan, K.M., Martin, W.K., Andersen, E.H., Williams Avram, S.K., Johns, J.M. & Robinson, D.L. (2017) Use of fast-scan cyclic voltammetry to assess phasic dopamine release in rat models of early postpartum maternal behavior and neglect. Behav Pharmacol 28, 648–660.

Sichel, D.A., Cohen, L.S., Robertson, L.M., Ruttenberg, A. & Rosenbaum, J.F. (1995) Prophylactic estrogen in recurrent postpartum affective disorder. Biological Psychiatry 38, 814–818.

Siegel, H.I. & Rosenblatt, J.S. (1975) Estrogen-induced maternal behavior in hysterectomized-overiectomized virgin rats. Physiol Behav 14, 465–471.

Stoffel, E.C. & Craft, R.M. (2004) Ovarian hormone withdrawal-induced “depression” in female rats. Physiol Behav 83, 505–513.

Suda, S., Segi-Nishida, E., Newton, S.S. & Duman, R.S. (2008) A postpartum model in rat: behavioral and gene expression changes induced by ovarian steroid deprivation. Biol Psychiatry 64, 311–319.

Winstanley, C.A., LaPlant, Q., Theobald, D.E.H., Green, T.A., Bachtell, R.K., Perrotti, L.I., DiLeone, R.J., Russo, S.J., Garth, W.J., Self, D.W. & Nestler, E.J. (2007) DeltaFosB induction in orbitofrontal cortex mediates tolerance to cocaine-induced cognitive dysfunction. J Neurosci 27, 10497–10507.

Yang, R., Zhang, B., Chen, T., Zhang, S. & Chen, L. (2017) Postpartum estrogen withdrawal impairs GABAergic inhibition and LTD induction in basolateral amygdala complex via down-regulation of GPR30. Eur Neuropsychopharmacol 27, 759–772.

Zachariou, V., Bolanos, C.A., Selley, D.E., Theobald, D., Cassidy, M.P., Kelz, M.B., Shaw-Lutchman, T., Berton, O., Sim-Selley, L.J., Dileone, R.J., Kumar, A. & Nestler, E.J. (2006) An essential role for DeltaFosB in the nucleus accumbens in morphine action. Nat Neurosci 9, 205–211.

Zhang, S., Hong, J., Zhang, T., Wu, J. & Chen, L. (2017) Activation of Sigma-1 Receptor Alleviates Postpartum Estrogen Withdrawal-Induced “Depression” Through Restoring Hippocampal nNOS-NO-CREB Activities in Mice. Mol Neurobiol 54, 3017–3030.

Zhang, Z., Hong, J., Zhang, S., Zhang, T., Sha, S., Yang, R., Qian, Y. & Chen, L. (2016) Postpartum estrogen withdrawal impairs hippocampal neurogenesis and causes depression- and anxiety-like behaviors in mice. Psychoneuroendocrinology 66, 138–149.

